# SpatioMark: Quantifying the impact of spatial proximity on cell phenotype

**DOI:** 10.1101/2024.12.04.626887

**Authors:** Sourish S Iyengar, Alex R Qin, Nicholas Robertson, Andrew N Harman, Ellis Patrick

## Abstract

As research advances in spatially resolving the biological archetype of various diseases, technologies that capture the spatial relationships between cells are demonstrating increasing value. Whilst there are an increasing number of analytical methods being developed to identify the complex web of interactions between cells, the downstream impacts of these cell-cell relationships are under explored. Here, we present SpatioMark, a statistical framework that simplifies the assessment of gene or protein expression in relation to the spatial proximity of different cell types. We demonstrate its performance across spatial proteomics and transcriptomics datasets and link identified relationships with differences in patient survival. We highlight key challenges in identifying changes in molecular markers associated with the localisation of cells and propose corrections which reduce artefact induced relationships. SpatioMark is implemented in the Statial R package hosted on the Bioconductor Project, ensuring interoperability with existing spatial analysis tools. Ultimately, this work highlights strategies for identifying and interpreting changes in cell phenotype associated with cellular relationships in spatial omics data, with broad applicability across various multiplexed imaging platforms.

## Introduction

Interactions between cells are important for homeostasis as well as a primary driver of biological change in healthy development and disease. Identifying key cellular interactions and their contributions to biological processes is therefore important for developing therapeutics and diagnostics (Armingol et al., 2021). To study cellular interactions, powerful spatial imaging technologies have been developed that are capable of multiplexed measurements of RNA and/or protein abundance at spatial and single cell resolutions *in situ* (Lewis et al., 2021). These highplex imaging technologies provide the capacity to visualise progressive changes in a cell’s transcriptome or proteome as spatial proximity changes between cells, thereby providing an avenue for identifying molecular changes associated with cellular interactions.

Identifying spatial relationships that drive key phenotypic changes in cells is a challenging task with numerous proposed methodological approaches. For example, recent methods use both cell marker and cell type data to identify cell-cell interactions by analysing co-expressed ligand-receptor pairs (Cang et al., 2023; Dries et al., 2021); however, this approach relies on those measured pairs being present in the chosen ligand-receptor database and so may overlook more subtle or novel interactions that can influence a cell’s molecular profile. Alternatively, other approaches can identify general expression changes in cells associated with their environments, however these methods either do not identify the specific interactions between cell types associated with these changes (Arnol et al., 2019; Tanevski et al., 2022), or identify changes associated with the composition of cells in a spatial domain, niche or region of tissue (Mason et al., 2024). Thus, there still remains a need for spatial methods that can flexibly detect both well-characterised and novel cellular interactions while providing a clear link between specific cell types and their spatial context within the tissue microenvironment.

Here we introduce SpatioMark, a statistical framework for identifying the effects of cell-cell proximity on the molecular profiles of cells measured by cell-resolution spatial omics technologies. This framework is implemented in the Statial R package hosted on the Bioconductor Project, and provides users with easily accessible functionality that is interoperable with other spatial analysis packages on Bioconductor. We demonstrate that while identifying changes in marker expression associated with the proximity between cell types (cell-cell-marker relationships; CCM) can be proposed as a conceptually straightforward hypothesis test, there are complexities to how these tests can be applied and interpreted. Irrespective of these complications, using cell-resolution MIBI-TOF, CODEX and Xenium spatial omics data we will demonstrate how SpatioMark can be used to identify CCM associated with patient survival in cancer cohorts.

## Methods

### Evaluation data

A co-detection by indexing (CODEX) dataset which aimed to characterise the immune tumour microenvironment in advanced-stage colorectal cancer (Schurch et al., 2020) was used to evaluate the behaviour of SpatioMark. The dataset consists of 35 advanced colorectal cancer patients, with 4 images per patient for a total of 140 images. Each image is marked with a 56-antibody panel to characterise a total of 24 distinct tumour and immune cell populations. For the purpose of exploring transitioning cell states with SpatioMark, cell types which reflected transitioning periods in certain cell types were simplified into 18 overarching cell types (**Supplementary Table 1**). Overall, a dataset consisting of 240k segmented cells was downloaded from Schurch et al., along with clinical information including patient tumour grade, tumour type, and patient survival.

A multiplexed ion beam imaging by time of flight (MIBI-TOF) dataset aimed at characterising the tumour microenvironment in triple negative breast cancer (Keren et al., 2018) was used to evaluate the predictive capabilities of features generated by SpatioMark. The Keren et al. dataset consisted of 38 patients with one image each. Each image was marked by a 36 antibody panel with 17 distinct cell types identified by the authors. Overall, a total of 197k segmented cells were downloaded from Keren et al., along with clinical information on patient survival.

A single multiplexed breast cancer image obtained using the 10x Genomics Xenium platform was used to evaluate the effect of different segmentation techniques on the performance of lateral spillover correction in SpatioMark. Data was obtained from a manuscript that compared multiple segmentation approaches (Fu et al., 2024) and comprised 103k segmented cells with 18 distinct cell types.

### Measurements of cell colocalization

SpatioMark is a spatial regression technique. In the package we have implemented two spatial metrics which can be used to quantify the spatial associations between cells.

#### Abundance

Cell abundance was used to quantify the number of each cell type that surrounds each individual cell. We define the abundance measure, *n_i,B_*(*r*), as the number of cells of cell type B within a *r* micron radius of the cell *i*.

#### Distance

We define *d_i,B_*(*r*) as the distance of the closest cell type B of the cell *i* in a *r* micron radius of cell *i.* To reduce the impact of high leverage cells, we constrain the maximum value of this distance. A maximum ceiling value of 200 was used as the defaults for the analysis.

### Modelling cell-interactions

Changes in a cell type’s state are proxied through variations in a cell’s marker expression. A cell’s state was modelled as a function of abundance or proximity to another cell type using ordinary least squares (OLS) regression:

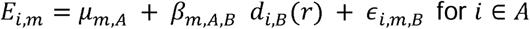

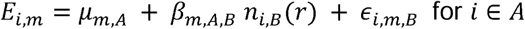

Where *E_i,m_* is the expression level for marker *m* in cell *i*, *µ_m,A_* is the average expression of marker m for cell type *A* and *d_i,B_*(*r*) is the distance of the closest cell type *B* within a *r* micron radius of the cell *i*. For abundance, *n_i,B_*(*r*) is the number of cells of cell type *B* within a r micron radius of a cell *i*. *β_m,A,B_* is the regression coefficient of how marker m changes in cell type A with respect to *d_i,B_*(*r*) or *n_i,B_*(*r*). Finally, *∈_i,AB,m_* is a random error term for the expression of marker m in cell *i* with proximity to cell type *B*.

To identify CCM relationships that are consistent across multiple images, models were fit individually for each image, pairwise cell type and marker combination. Relationships with less than 100 cells were excluded. Coefficients across images were averaged per patient. The statistical significance of pairwise cell type and marker combination relationships was quantified by applying a two sided t-test across the regression coefficients to test if their mean was zero. The false positive rate was controlled by a Bonferroni correction.

### Correcting for marker contamination

Contamination was assessed by fitting a random forest model (from the ranger R package) using all images to predict a cell’s type given its marker expression values. The purity scores of a cell were defined as the predicted probability of the cell belonging to each cell type. Let (*X_1_*, *X_2_*,. .. , *X_m_* ) represent the marker expression values, and let (*Y_1_*, *Y_2_* , . . ., *Y_n_*) represent the cell type of each cell.

1. Train a Random Forest model to predict the cell type labels (*Y_1_*, *Y_2_*,. .., *Y_n_)* for each cell, given the marker expression matrix (*X_1_*, *X_2_*,. .., *X_m_*).
2. Use the trained model to predict the probability for each cell type *t* for each cell.

The probability scores for each cell type were used as additional model covariates in equations to attempt to mitigate the effect of marker contamination on the biological validity of statistical inferences. The intended effect of this is to reduce the variation in a cell type’s marker expression explained by the distance/abundance to another cell type when marker spillover is driving the marker expression change. These probabilities were factored in as covariates into our linear equation such that

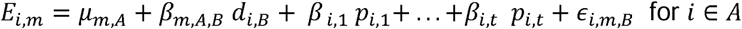

Where *p_i,n_* is the probability of each cell *i* belonging to each of the *t* cell types as predicted by the random forest model.

The effectiveness of these contamination correction covariates was evaluated by comparing the number of interactions involving cell state markers relative to cell type markers in the top 100 false positives (cell type markers) ranked by statistical significance. Cell type markers were defined as markers that are expected to only appear in the cell type the marker they are specific to (**Supplementary Table 2**). A partial ROC curve was constructed by identifying the most significant (sorted by ascending p-value) cell state relationships and cell type relationships with and without contamination correction and progressively adding the modelled interaction with the highest model significance until the first 100 false positives were reached.

The robustness of the contamination correction approach on alternative imaging modalities and segmentation quality was also assessed using spatial transcriptomics data. An image of varying segmentation quality was applied by using Voronoi, BIDCell and Nuclei segmentation algorithms. The Voronoi algorithm is expected to have the highest level of marker spillover given its less conservative nature. While the Nuclei segmentation is expected to have the least spill over because of its conservative nature at the trade off capturing less marker expression information. The BIDcell algorithm is expected to form a middle ground between capturing sufficient marker expression information and avoiding excessive marker contamination. The linear models were applied with and without contamination correction method across these varying segmentation methods and performance was evaluated using a partial ROC curve as before.

### Modelling the relationship between cell interactions and patient outcomes

Linear model test-statistics were extracted for each relationship fitted with a radius of 200 microns and used as modelling covariates to predict patient survival. To ensure biologically relevance, relationships which involved fewer than 20 cells in a particular image were removed. Following, relationships with greater than 5% missingness across all images were filtered out prior to fitting univariate Cox proportional hazards survival models. Relationships with a Bonferroni adjusted p-value < 0.05 were considered significant.

Patient survival was also modelled using survival random forest models implementation with the ranger package (Wright & Ziegler, 2017) and evaluated with 20 repeats of 3-fold cross-validation using the ClassifyR package (Strbenac et al., 2015). In the training folds of cross-validation, test-statistics for each relationship were ranked by their significance with survival in univariate Cox proportional hazards models models. The top 10 most significant relationships were used to build survival random forest models. The performance of these models were evaluated by calculating the concordance index of model predictions on the test folds. The concordance index was calculated as:

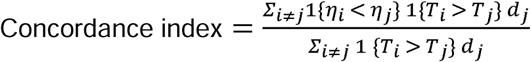

Where 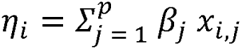 is the estimated risk score for patient *i* given their characteristics and *T_i_* is the length of time patient *i* has survived for as of their last status record. *d_i_ ∈* [0,1] is the binary indicator of the comparison patient *j* censoring whether the patient is alive or dead based on their last known record.

## Results

Here we present SpatioMark, a framework for quantifying changes in protein or transcript marker expression associated with proximity between cell types (cell-cell-marker (CCM) relationships). Cells can influence nearby cells through a variety of secreted factors and direct cell-to-cell contact. As cells become closer in spatial proximity, certain cell types are capable of influencing the marker expression of other cell types (**Figure 1A**). To identify the relevance and magnitude of these CCM relationships, SpatioMark first quantifies spatial proximity of one cell type to another using either a distance or cell abundance derived spatial proximity metric (**Figure 1Bi**), denoises for segmentation artefacts (**Figure 1Bii**), then fits a linear model between the measure of spatial proximity and the expression of a cell marker (**Figure 1Biii**). In the following we will demonstrate this functionality and that the identified CCMs can vary across patient cohorts.

**Figure 1:**
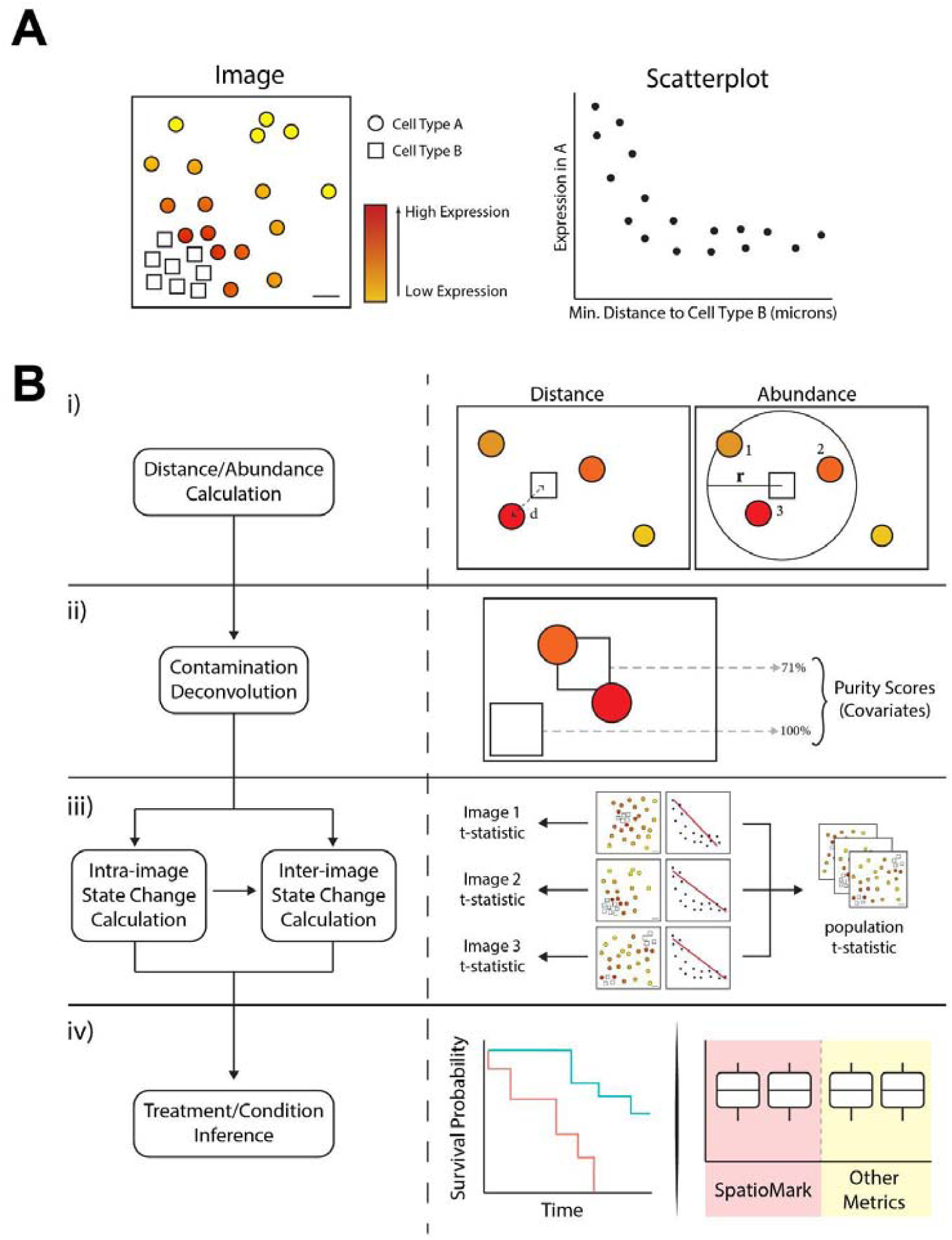
The SpatioMark framework. **(A)** An illustrative example of spatial proximity between cell type A and cell type B being associated with marker expression in cell type A. **(B)** SpatioMark workflow (i) Spatial proximity metric calculation for either distance or abundance (ii) Deconvolution schematic demonstrating the calculation of purity scores of each cell type followed by factoring into our linear model as covariates (iii) fitting of linear model (iv) downstream analysis using SpatioMark outputs as metrics to segregate patients by survival.

To demonstrate SpatioMark’s ability to extract relevant CCMs, we applied the SpatioMark framework to spatial proteomic images of colorectal cancer (Schürch et al., 2020). The Schürch dataset contains 140 images obtained from 35 patients with cells annotated into 29 discrete cell types. For the purpose of downstream processing SpatioMark we’ve simplified this into 18 cell types, combining discrete annotations of transitioning cell states such as combining CD163+ and CD163-macrophages into macrophages. Using the minimum distance between cell types as a proximity metric, the SpatioMark framework identifies 401 CCMs with a Bonferroni adjusted p-value less than 0.05 and a mean coefficient greater than 0.01 (**Figure 2A**). The expression of CD163 by macrophages changes with macrophage proximity to tumour cells has a small adjusted p-value (7.7 x 10^-5^) and large coefficient (0.061; 99th percentile of all significant CCMs) relative to other CCMs (**Figure 2A**). A representative image of this relationship demonstrates that average CD163 abundance in macrophages is lowest when macrophages are close to tumour cells (**Figure 2B**). Evidence of this Tumour-Macrophage-CD163 CCM is observed when using both minimum distance and abundance as metrics to quantify cell proximity (**Figure 2C, Figure 2D**), consistent with the strong concordance observed across all relationships for the two spatial metrics (**Figure 2E**).

**Figure 2:**
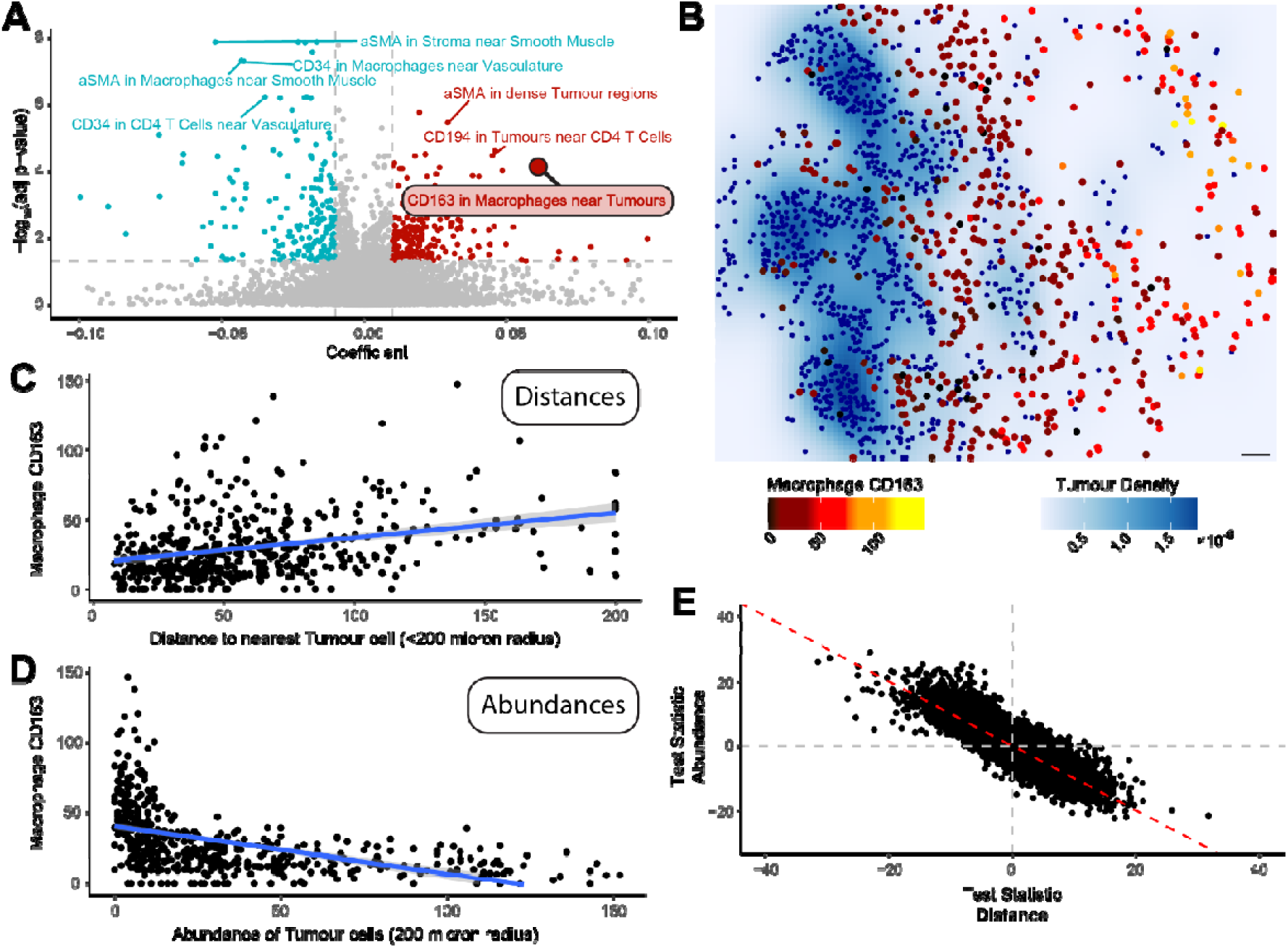
Application of SpatioMark to a colorectal cancer CODEX dataset. The SpatioMark framework was used to identify cell-cell relationships and their effects on protein markers in colorectal cancer. **(A)** Volcano plot of relationships comparing coefficient against the -log10 adjusted p-values. Positive relationships above an adjusted p-value of 0.05 and with a greater coefficient than |0.006| are highlighted in red, and negative relationships are highlighted in blue. **(B)** Image 5A from Schürch et al. (2020); Tumour cells are highlighted in blue, macrophages highlighted on a gradient from black to yellow. Density of tumour cells are plotted on a white to dark blue scale. **(C)** Scatterplot showing the expression of CD163 in macrophages against the distance metric between macrophages and tumour cells. Blue line represents the linear model. **(D)** Scatterplot showing the expression of CD163 in macrophages against the abundance metric of tumour cells within a 200 micron radius of macrophages. Blue line represents the linear model. **(E)** Scatterplot of test statistics for abundance against the test statistics for distance.

Lateral marker spillover is contributing to many of the identified CCM relationships. This occurs when markers from one cell are mistakenly attributed to nearby cells due to segmentation errors, interleaving of cell membranes of adjacent cells, or technical artefacts. Among the top-ranked CCMs, CD34, a marker predominantly expressed in vascular cells, shows increased expression in macrophages located near vascular cells (adjusted p = 1.7x 10^-23^, **Figure 3A**, **Figure 3B**). Similarly, aSMA, a smooth muscle marker, was found to increase in macrophages near smooth muscle cells (adjusted p = 9.4 x 10^-21^). These markers are highly cell-type specific and should not be present in macrophages under normal biological conditions. Acknowledging that marker expression in a cell might be contaminated by nearby cells, we estimated the probability of each macrophage cell being a vasculature cell. That is, a quantification of how similar the expression profile of a cell is to the average expression profile of its cell type relative to other cell types. In this case, a quantification of how similar each macrophage cell is to a vasculature cell or a smooth muscle cell. Using this metric, the increase of CD34 in macrophages near vascular cells and aSMA in macrophages near smooth muscle appears to be driven by contamination and low purity (**Figure 3C**, **Figure 3D**), indicating that spillover is responsible for the observed CCM. These spillover-induced relationships highlight the importance of interpreting relationships in spatial omics data with an understanding that they may be induced by technical artefacts.

**Figure 3:**
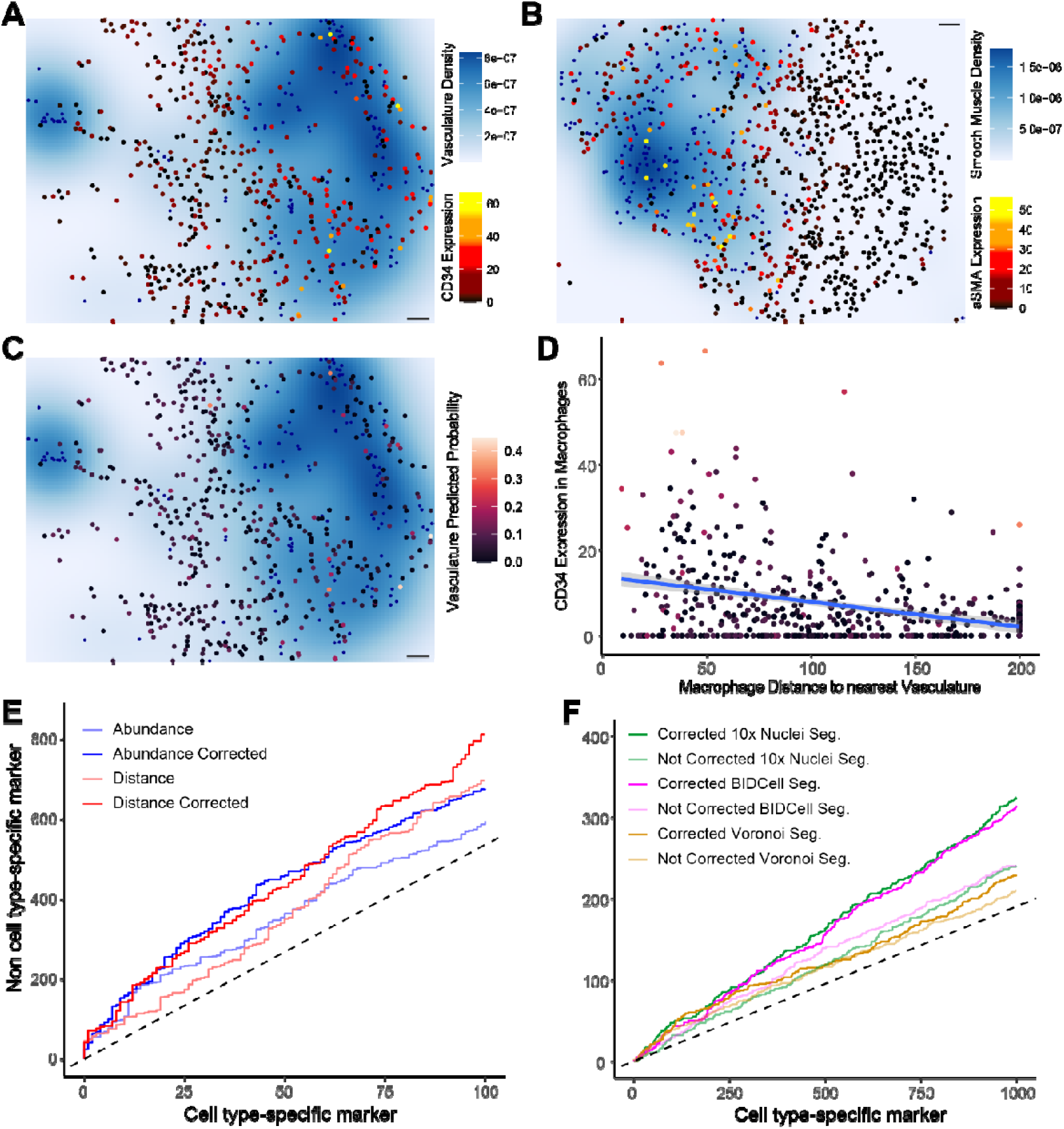
Effects of lateral cell spillover correction on cell-cell-marker relationships in SpatioMark. **(A)** Image 65A from Schürch et al. (2020); B cells are highlighted in blue, CD11b+ monocytes are highlighted on a gradient from black to yellow. Density of tumour cells are plotted on a white to dark blue scale. **(B)** Scatterplot showing the expression of CD20 in CD11b+ monocytes against the distance metric between B cells and CD11b+ monocytes. Blue line represents the linear model. **(C)** Image 65A from Schürch et al., (2020); predicted probability of a cell classified as CD11b+ monocyte being designated as a CD11b+ monocyte is used as a proxy measure of purity. **(D)** Scatterplot of correlation between test statistics for all cell-cell-marker relationships corrected for lateral marker spill over against uncorrected test statistics. **(E)** Partial ROC curve displaying true cell state markers on the y-axis compared to the false cell type markers on the x-axis, showing the trade-off between detecting relationships involving cell state and cell type markers for different metrics and corrections. **(F)** Partial ROC curve displaying true cell state markers on the y-axis compared to the false cell type markers on the x-axis, for different segmentation techniques, with and without lateral spillover correction.

To reduce the impact of lateral marker spillover on identified CCMs, we implemented a spatial multiple regression model that included cell type probability scores. Cell type probability scores were estimated by a random forest model that predicted the likelihood of each cell belonging to a specific cell type based on its marker expression profile. To evaluate the effectiveness of this correction, we compared the ranks of cell type-specific markers such as CD4, which should not be identified as CCMs, with other markers, treating cell type-specific markers as false positives. The uncorrected models for both distance and abundance metrics identified a lower number of true positive non-cell type-specific CCMs (distance - 697 TPs; abundance - 587 TPs), while the corrected model increased the number of true-positive CCMs (distance - 812 TPs; abundance - 675 TPs) by 16.5% and 15% respectively for the first 100 false positives (**Figure 3E**). Although not perfect, this correction method does reduce the number of false-positive relationships caused by spillover.

The issue of marker spillover is not limited to spatial proteomics assays. We applied SpatioMark to a breast cancer Xenium dataset and again observed cell type-specific CCMs, which should not occur, appearing in the analysis (**Figure 3F**). This problem was present across multiple segmentation methods. Specifically, we tested three segmentation techniques: Voronoi, a non-conservative approach where cells are defined based on proximity and are prone to high levels of marker spillover; Nuclei segmentation, a more conservative method that isolates cells based on nuclear boundaries, which reduces spillover but may lose important marker expression information; and BIDCell segmentation, a deep learning-based method that aims to balance capturing sufficient marker information while minimising spillover. BIDCell had the best performance for uncorrected tests. The Voronoi segmentation, as expected, resulted in the highest degree of spillover induced CCMs due to its liberal approach to defining cell boundaries. Lateral spillover correction improved the performance of the Nuclei segmentation and BIDcell segmentation, but did not have a large impact on the performance of the Voronoi segmented data. These differences in performance emphasise the need for high quality segmentation methods and the limitations of accounting for artefacts post-segmentation.

To explore whether the features of SpatioMark could be used beyond interrogating cellular relationships, we investigated if there were CCMs that are associated with patient survival. To achieve this we used the SpatioMark statistics as explanatory variables in Cox proportional hazards models. We found 7 SpatioMark features which were predictive of patient survival with a Bonferroni adjusted p-value of less than 0.05 (**Figure 4A**). The two relationships with the largest coefficient in each direction were “CD45RA in Tumours near Smooth Muscle” and “Na+/K+ ATPase in Tumours near Macrophages”. We found that increased expression of CD45RA in Tumours near Smooth Muscle cells were correlated with reduced patient survival, whilst increased expression of Na+/K+ ATPase (NKA) in tumours near macrophages was positively correlated with patient survival (**Figure 4B**). In isolation only the expression of Na+/K+ ATPase in Tumours was significantly associated with survival, whilst other markers in their respective cell types and the proportion of these cell types were not significantly associated with survival (**Supplementary Table 3**). These results highlight the value of examining cellular relationships from multiple perspectives.

**Figure 4:**
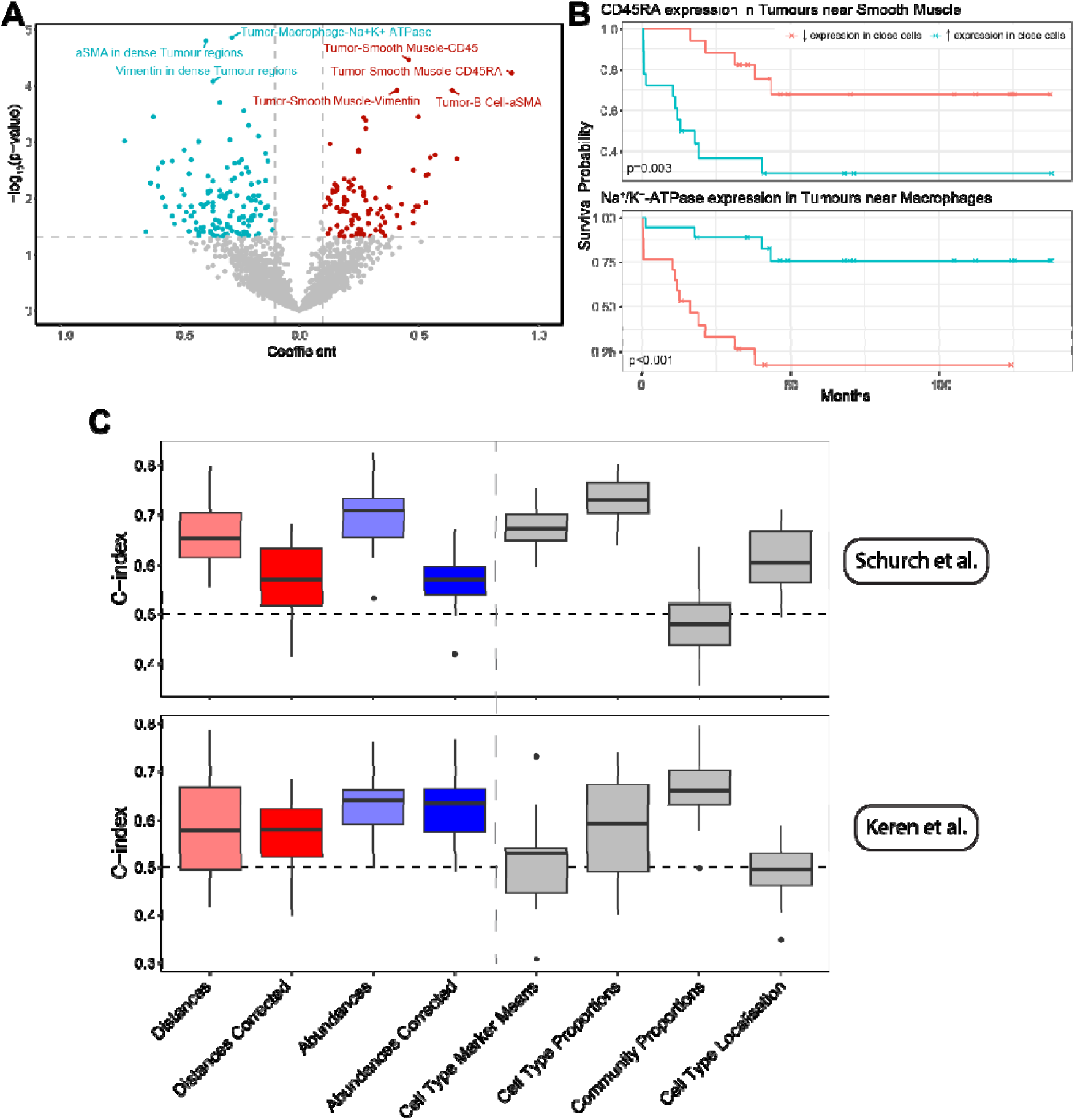
Survival analysis and classification using SpatioMark. **(A)** Volcano plot of relationships comparing coefficient against the -log10 adjusted p-values. Positive relationships above an adjusted p-value of 0.05 and with a greater coefficient than |0.01| are highlighted in red, and negative relationships are highlighted in blue. **(B)** Kaplan Meier curves of the relationship between CD5 expression in CD8 T cells and proximity to Tumour cells. The curves are stratified by the median value of each method. Reported p-values are produced from a cox proportional hazard model fitted on the original features values prior to splitting them on the median. **(C)** Survival classification results for 2 datasets were produced using regularised Cox Regression models trained on the four SpatioMark outputs and other spatial/non-spatial features. The models were evaluated with cross-validation and assessed using Harrell’s C-index (ranges from 0 to 1), where a higher C-index indicates better performance in survival classification.

Finally, we aimed to contextualise the strength of the relationship between SpatioMark features with patient survival relative to other quantifications. To do this we compared survival models using SpatioMark features to multiple feature sets; the proportions of the cells types in each image; proportion of different spatial communities identified by lisaClust (Patrick et al., 2023); and pairwise distances between cell types generated by spicyR (Canete et al., 2022). We made these comparisons in the Schürch colorectal cancer dataset and a triple negative breast cancer dataset assayed with MIBI-TOF (Keren et al., 2018). For the Schürch data SpatioMark features were the most predictive of survival producing the highest C-Index (**Figure 4C**). For both of the datasets, correcting for contamination reduced predictive performance indicating that while spillover contamination impacts interpretation of CCMs, it is still informative. In the Schürch data, using minimum distances to quantify spatial relationships had superior performance while the abundance metric had superior performance in the Keren data. While we do not expect SpatioMark features to perform competitively in all datasets, these results do support the value of assessing a variety of metrics in spatial omics data when searching for associations with survival.

## Discussion

In this paper we presented SpatioMark, a framework for identifying changes in cell phenotypes related to spatial proximity of cells. This framework is implemented in the Statial package hosted on Bioconductor, providing accessible functionality to facilitate the testing of a well defined hypothesis - does a phenotypic marker on a cell type change when that cell type is near another cell type? When demonstrating the CCM relationships that SpatioMark can identify in a spatial proteomics dataset, we highlighted a clear limitation for interpreting identified CCMs caused by lateral marker spillover artefacts. We showed that we could partially correct for this phenomenon post-segmentation in both spatial proteomics and spatial transcriptomics datasets. Finally, we demonstrated that CCMs can be associated with patient survival in two cancer datasets and were informative features to include in predictive models. The ability of SpatioMark to identify such biologically relevant CCMs highlights its value in exploring cell-cell interactions and their impact on clinical outcomes. This framework provides a powerful tool for uncovering novel relationships in spatial omics datasets that will contribute to our growing understanding of the tumour microenvironment and other systems.

One of the key strengths of SpatioMark is its ability to summarise changes in CCM relationships across multiple images, offering biologically relevant insights. Among the significant findings identified by SpatioMark is the expression of CD163 by macrophages near tumours. CD163, an anti-inflammatory marker, is typically downregulated as macrophages shift to a pro-inflammatory state in response to tumour cells (Etzerodt & Moestrup, 2013). This observation aligns with existing literature, such as a study in colorectal cancer that reported lower CD163 expression in macrophages near tumours, which is linked to immune cell recruitment to target and destroy cancer cells (Shabo et al., 2014). Thus, SpatioMark can detect biologically plausible relationships, revealing novel interactions between cells and their effects on molecular expression.

Beyond identifying interactions, SpatioMark can also link CCMs to clinically important metrics such as patient survival. For example, the analysis revealed that increased CD45RA expression in Tumours near smooth muscle cells was associated with poorer patient survival. It is highly likely that this is a relationship caused by lateral spill-over, as CD45RA is a highly specific immune marker of memory T cells. In effect, SpatioMark is picking up clinically relevant associations between 3 cell types, tumour cells, smooth muscle cells, and memory T cells, where their collective interaction leads to poorer patient survival. By linking these molecular interactions to patient outcomes, SpatioMark extends our understanding of how immune suppression mechanisms in the tumour microenvironment impact survival. Additionally, SpatioMark uncovered relationships involving Na+/K+ ATPase (NKA) in tumours near macrophages, which has been shown to either promote or inhibit cellular proliferation depending on cell type (Bejček et al., 2021; Khajah et al., 2018; Prassas et al., 2011). This suggests a complex interplay between tumour cells and macrophages, with potential implications for tumour growth and immune evasion.

Imperfect segmentation in spatial datasets, leading to lateral marker spillover, often results in the misattribution of marker expression between neighbouring cells (Bai et al., 2021; Berg et al., 2019). SpatioMark addresses this issue by incorporating post-segmentation corrections, using cell type probability scores from cell type deconvolution, to reduce false-positive CCM relationships. Lateral marker spillover can be reduced by advanced segmentation approaches like BIDcell (Fu et al., 2024) that are optimised to increase purity of segmented cells or through post-segmentation **lateral spillover compensation methods** like REDSEA (Bai et al., 2021) that correct for spillover across cell borders and within the cell body. While upstream segmentation techniques are evolving, corrections like those in SpatioMark are essential to ensure reliable insights from spatial data.

One key limitation of our approach is the use of ordinary least squares regression, which may not fully capture the non-linear and heteroscedastic nature of relationships in spatial omics data. The residuals of the fitted models often deviated from normality due to zero marker expression, and the variance in marker expression was often not constant across predictors. Although we explored more flexible models, such as generalised additive models and polynomial regression, these approaches did not significantly alter our findings, with all approaches identifying an overwhelming number of relationships with small p-values. Despite its limitations, linear regression offers a practical balance between interpretability and accuracy, future research should focus on developing more sophisticated models that better capture the complexity of the data while maintaining interpretability.

## Data & Code Availability

All datasets used in this study are publicly available. The CODEX advanced colorectal cancer and MIBI-TOF triple negative dataset are available in the SpatialDatasets bioconductor package (Ameen et al., 2023). Code used to generate the figures in this manuscript can be found on github: https://github.com/SydneyBioX/SpatioMark. SpatioMark is available in the Statial R package found on bioconductor: https://bioconductor.org/packages/release/bioc/html/Statial.html.

## Author Contributions

All authors contributed to the interpretation of results and writing of the manuscript. EP and SSI conceived the project with SSI and ARQ performing the main analysis. SSI, ARQ and NR contributed to the design and implementation of the R package and functionality.

## Acknowledgements

We thank the scientific community for proactively making their data and code publicly available. This work has been partly supported by The University of Sydney and an Australian Research Council Discovery Early Career Researcher Award (DE200100944) funded by the Australian Government.

## Supplementary Material

**Supplementary Table 1:** Transitionary cell types in the original Schurch et al. manuscript which have been collapsed into a single cell type.

| Initial cell type | New cell type |
| --- | --- |
| CD163+ Macrophages | Macrophages |
| CD68+ GranzymeB+ Macrophages | Macrophages |
| CD11b+ CD68+ Macrophages | Macrophages |
| CD68+ Macrophages | Macrophages |
| CD68+ CD163+ Macrophages | Macrophages |
| CD45RO+ CD4 T cells | CD4 T cells |
| GATA3+ CD4 T cells | CD4 T cells |
| CD4 T cells | CD4 T cells |

**Supplementary Table 2:**
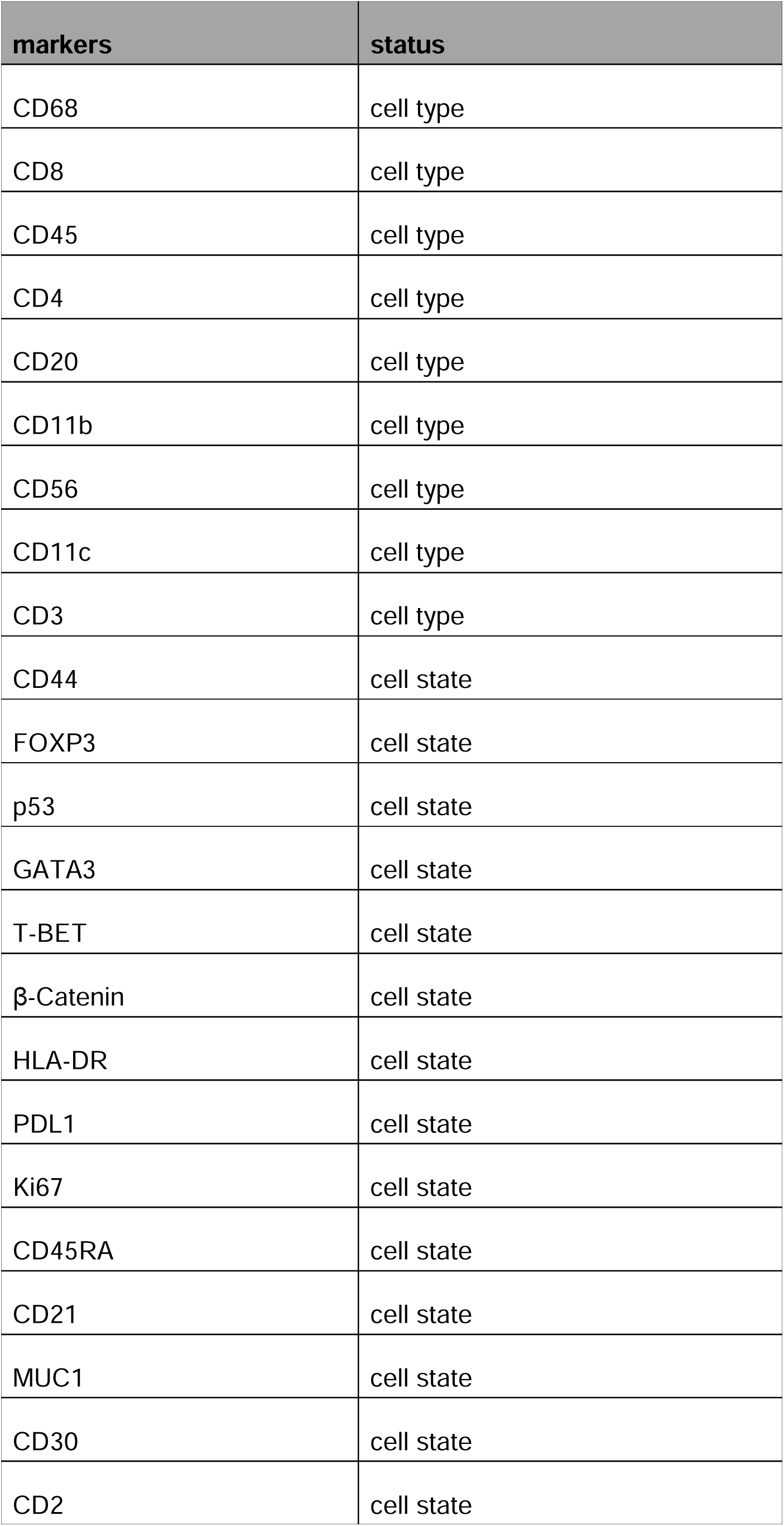
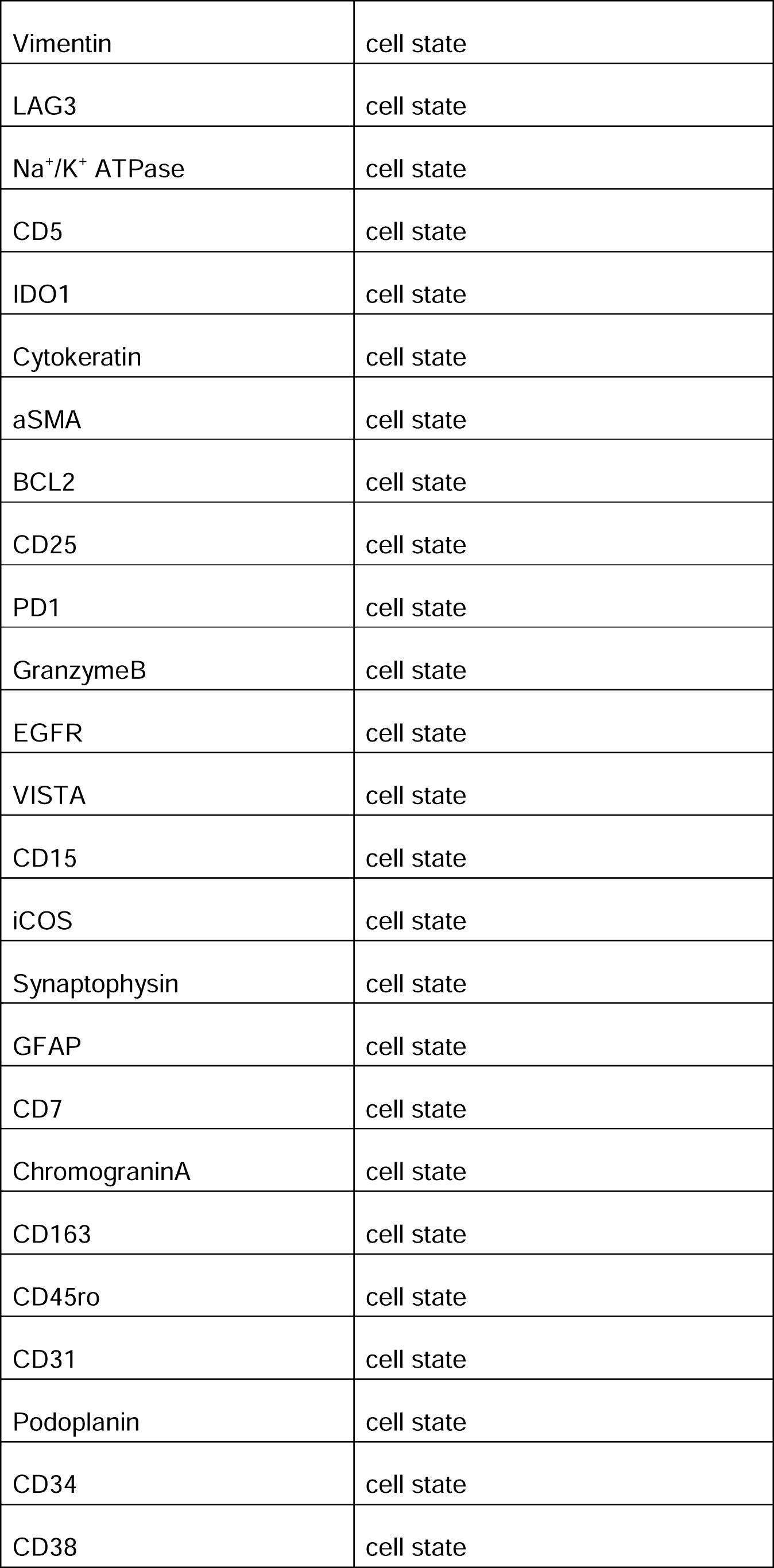
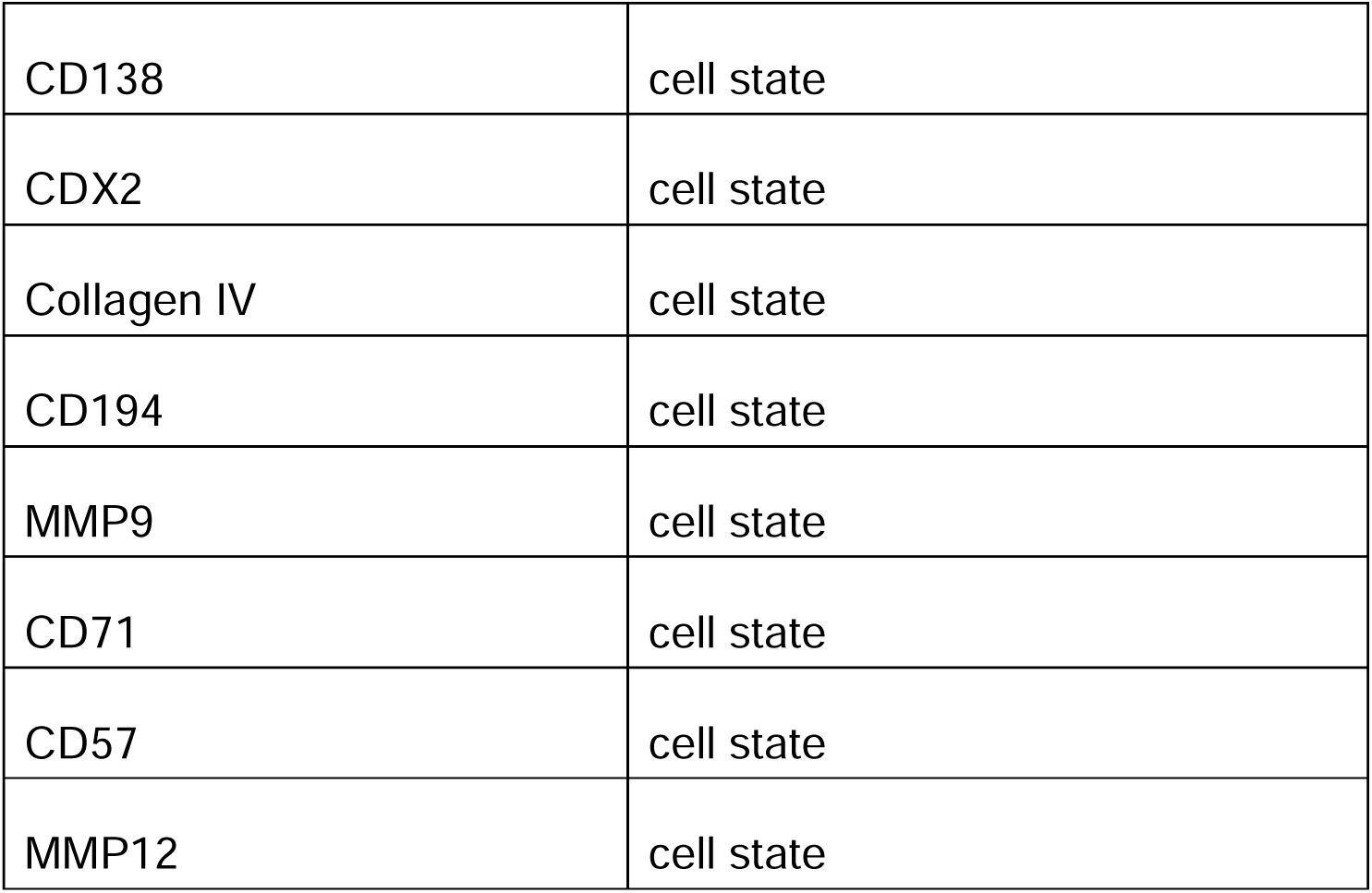
Designations of cell type vs cell state used for partial ROC curve calculations in Figure 3.

| markers | status |
| --- | --- |
| CD68 | cell type |
| CD8 | cell type |
| CD45 | cell type |
| CD4 | cell type |
| CD20 | cell type |
| CD11b | cell type |
| CD56 | cell type |
| CD11c | cell type |
| CD3 | cell type |
| CD44 | cell state |
| FOXP3 | cell state |
| p53 | cell state |
| GATA3 | cell state |
| T-BET | cell state |
| β-Catenin | cell state |
| HLA-DR | cell state |
| PDL1 | cell state |
| Ki67 | cell state |
| CD45RA | cell state |
| CD21 | cell state |
| MUC1 | cell state |
| CD30 | cell state |
| CD2 | cell state |
| Vimentin | cell state |
| LAG3 | cell state |
| Na<sup>+</sup>/K<sup>+</sup> ATPase | cell state |
| CD5 | cell state |
| IDO1 | cell state |
| Cytokeratin | cell state |
| aSMA | cell state |
| BCL2 | cell state |
| CD25 | cell state |
| PD1 | cell state |
| GranzymeB | cell state |
| EGFR | cell state |
| VISTA | cell state |
| CD15 | cell state |
| iCOS | cell state |
| Synaptophysin | cell state |
| GFAP | cell state |
| CD7 | cell state |
| ChromograninA | cell state |
| CD163 | cell state |
| CD45ro | cell state |
| CD31 | cell state |
| Podoplanin | cell state |
| CD34 | cell state |
| CD38 | cell state |
| CD138 | cell state |
| CDX2 | cell state |
| Collagen IV | cell state |
| CD194 | cell state |
| MMP9 | cell state |
| CD71 | cell state |
| CD57 | cell state |
| MMP12 | cell state |

**Supplementary Table 3:**
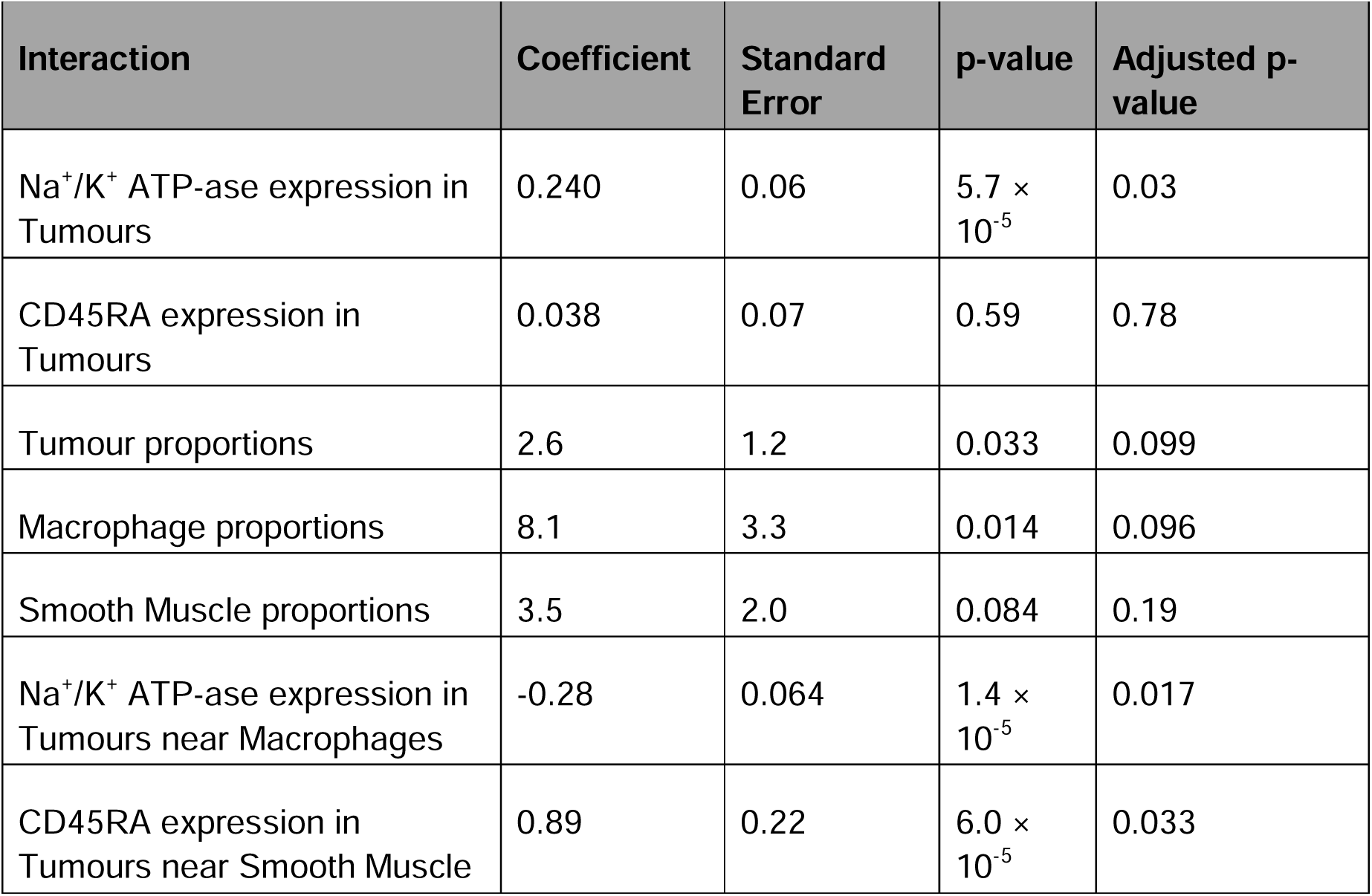
Survival model coefficients, standard errors, p-values, and adjusted p-values for different interaction types, from markers in cell types to cell type proportions and SpatioMark features.

